# Cell non-autonomous interactions during non-immune stromal progression in the breast tumor microenvironment

**DOI:** 10.1101/540112

**Authors:** Raditya Utama, Anja Bastian, Narayanan Sadagopan, Ying Jin, Eric Antoniou, Qing Gao, Yinghui J. Huang, Sailesh Gopalakrishna-Pillai, Peter P. Lee, Gurinder S. Atwal

## Abstract

**Summary:** The breast tumor microenvironment of primary and metastatic sites is a complex milieu of differing cell populations, consisting of tumor cells and the surrounding stroma. Despite recent progress in delineating the immune component of the stroma, the genomic expression landscape of the non-immune stroma (NIS) population and their role in mediating cancer progression and informing effective therapies are not well understood. Here we obtained 52 cell-sorted NIS and epithelial tissue samples across 37 patients from i) normal breast, ii) normal breast adjacent to primary tumor, iii) primary tumor, and iv) metastatic tumor sites. Deep RNA-seq revealed diverging gene expression profiles as the NIS evolves from normal to metastatic tumor tissue, with intra-patient normal-primary variation comparable to inter-patient variation. Significant expression changes between normal and adjacent normal tissue support the notion of a cancer field effect, but extended out to the NIS. Most differentially expressed protein-coding genes and lncRNAs were found to be associated with pattern formation, embryogenesis, and the epithelial-mesenchymal transition. We validated the protein expression changes of a novel candidate gene, C2orf88, by immunohistochemistry staining of representative tissues. Significant mutual information between epithelial ligand and NIS receptor gene expression, across primary and metastatic tissue, suggests a unidirectional model of molecular signaling between the two tissues. Furthermore, survival analyses of 827 luminal breast tumor samples demonstrated the predictive power of the NIS gene expression to inform clinical outcomes. Together, these results highlight the evolution of NIS gene expression in breast tumors and suggest novel therapeutic strategies targeting the microenvironment.

## 1 Introduction

Breast cancer is the second most prevalent type of cancer worldwide and accounts for nearly 25% of total cancer incidence in women [1]. Recent studies have highlighted the importance of the breast tumor microenvironment (TME) in mediating and regulating tumor progression with respect to the adaptive immune system and therapeutic intervention. TME contains not only cancer cells, but a significant fraction of non-cancer cells that collectively is termed stroma [2]. Importantly, high tumor stroma percentage (TSP) is associated with worse clinical outcome [2], including breast cancer [3]. Tumor stroma consists of multiple different cell types, including immune cells, cancer-associated fibroblasts, and endothelial cells. The non-immune stroma (NIS) is thought to play an important structural role in the tumor microenvironment, forming the connective tissues and regulating production of the extracellular matrix (ECM), while also providing functional support for tumor progression/regression via processes such as cell proliferation, immune evasion, angiogenesis, apoptosis, and metastasis promotion [4–6]. In the case of pancreatic tumors, dense NIS tissues have shown to be the main barrier for immune infiltration and have consequently become the biggest challenge in developing pancreatic cancer therapies [7]. However, the role of NIS in breast cancer is not well understood.

In this study, we hypothesized that changes in NIS gene activity track progression from normal breast tissue to primary tumor and metastatic tumor, reflecting functional molecular communication between cancer cells and their surrounding NIS. To test this, fresh primary and metastatic tumor samples from breast cancer patients were obtained and processed for RNA-seq analysis. In order to compare gene expression levels in noncancerous tissue, normal adjacent breast tissues from these patients and normal breast tissues from *BRCA* mutant (*BRCA*+) subjects undergoing prophylactic mastectomy were also collected. Live cells cultured from fresh tissue samples were sorted for NIS and epithelial/cancer cells followed by whole transcriptome bulk RNA-sequencing. A pseudoalignment method was used to map and quantify these transcripts onto a reference human genome. Prior to any genomic analyses, we performed rigorous computational quality control and batch correction to obviate technical artefacts in the RNA-seq data (see Methods).

Whole genome transcriptional patterns of NIS, when projected onto low dimensional representations, reveal a robust progression from normal to primary tumor to metastatic tumor sites. This observation suggests that, rather than acting as an inert structural background to the tumor, NIS dynamically interacts with the changing epithelial-derived cancer cells. Surprisingly, we observed that normal tissue adjacent to breast tumors have distinct expression profiles to noncancerous breast tissues from healthy patients – instead are closer to NIS from its adjacent primary tumor. Furthermore, NIS cells from primary tumors were transcriptionally distinct from metastatic tumors with significant changes in NIS gene activity. We investigated the Gene Ontology of significantly differentially expressed genes and found functional enrichment in pattern formation, embryogenesis, epithelial-mesenchymal transition (EMT), and immune response functions. Significant differentially expressed genes, exclusively expressed in NIS tissue, such as the *C2orf88* gene, were found to be downregulated in brain and liver metastatic sites as confirmed via IHC antibody validation. Further, we elucidated the NIS-epithelial crosstalk via ligand-receptor interactions and identified stronger ligand signaling originating from cancerous epithelial cells rather than from NIS cells. Together, these results support the concept of a cancer field effect [8, 9], whereby cancerous cells in the primary site exert their influence on the gene expression pattern of neighboring normal cells, as well as the crosstalk between NIS cells and cancer cells aiding in tumor progression

## 2 Results

### 2.1 Breast tissue samples and bulk RNA-seq

To investigate the transcriptional landscape of NIS and cancer cells, fresh breast tissue samples were obtained from patients at the City of Hope cancer center. Over a two year period, 79 samples from 52 patients were successfully acquired and analyzed in seven different batches. We classified the samples into four types: i) prophylactic normal breast tissue from women without breast cancer, ii) normal breast tissue ipsi-/contralateral to primary breast tumor, iii) primary tumor, and iv) metastatic tumor. By definition, prophylactic normal samples are tissues that were taken from tumor negative patients with *BRCA* mutations who have undergone a prophylactic mastectomy procedure. Ipsilateral normal samples were collected from macroscopically noncancerous tissue areas located adjacent to a primary tumor site and confirmed by a pathologist to be noncancerous. Contralateral samples were collected similarly to ipsilateral normal samples but from the opposite breast, which was prophylactically resected during the primary tumor surgery. Primary tumor samples were tumors growing at the primary breast site whereas metastatic samples were tumor tissues that metastasized from the primary tumor to different locations in the body, specifically brain, liver, lung, skin, and bone. In order to capture a representative coverage of every tissue, short-term (1 week) *in vitro* culturing procedures were implemented to expand the cells and isolate sufficient amounts of RNA. This step was essential due to the limited number of fibroblasts that could be isolated from each tissue sample.

We implemented a cell-sorting step, in order to isolate pure NIS cells that consisted primarily of cancer-associated fibroblasts (CAF) and endothelial cells from the tumor samples. In a previous study, our laboratory [10] showed that NIS could be generally identified by high expression of *CD44* but low expressions of *EPCAM* and *CD45*. Epithelial cells and cancer cells express high levels of *EPCAM* but low levels of *CD44* and *CD45*. By combining these three markers, we sorted cells from each tissue into NIS cells (*CD44*+, *EPCAM*-, *CD45*-) and epithelial/cancer cells (*EPCAM*+, *CD44*-, *CD45*-) via flow cytometry. Different cell populations found in NIS and epithelial samples are confirmed by expression of several canonical markers (Fig. 1b), such as epithelial (*EPCAM*, *CDH1*), immune (*CD45*, *CD3E*), and fibroblast-pericyte-endothelial (*FAP*, *RGS5*, *PECAM1*).

**Figure 1:**
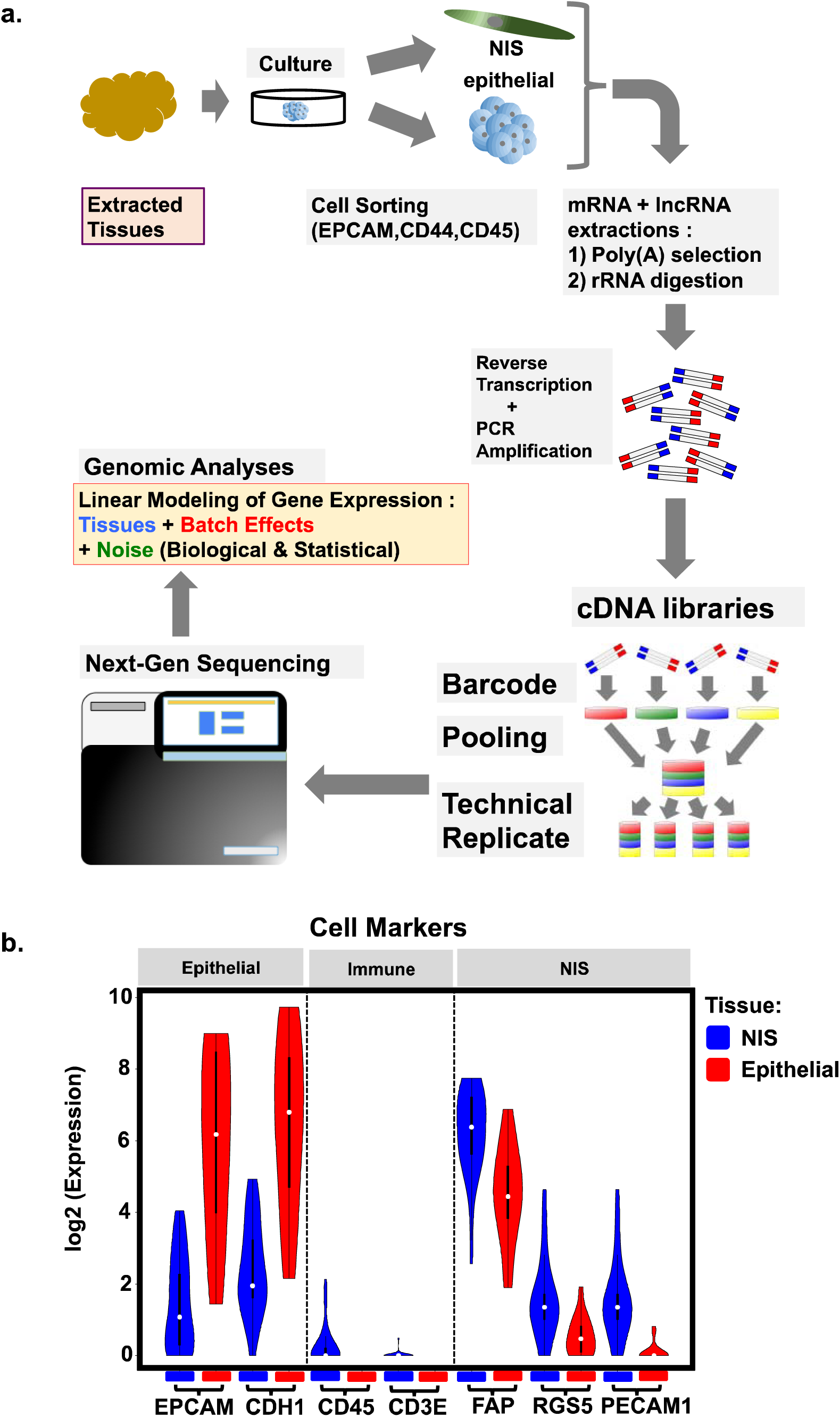
Framework of RNAseq protocol and analyses. **a)** Tissue samples were extracted from normal breast and luminal breast tumor biopsies. Cell markers were utilized to sort for non-immune stroma (NIS) (*EPCAM*-, *CD44*+, *CD45*-) and epithelial/cancer (*EPCAM*+, *CD44*-, *CD45*-). cDNA libraries pooled and randomized to obviate next-gen sequencing batch effects. **b)** Logfold expressions (tpm) of NIS samples (blue) and epithelial samples (red). Tissue cell markers for epithelial (*EPCAM*, *CDH1*), immune / T-cell (*CD45*, *CD3E*), and NIS / fibroblast-pericyte-endothelial (*FAP*, *RGS5*, *PECAM1*).

To extract messenger RNA (mRNA) and long noncoding RNA (lncRNA) from the entire RNA content produced by the samples, we excluded all significant contaminations such as rRNA, tRNA, and miRNA using standard library preparation methods. Once we acquired high yields of mRNA and lncRNA, sequencing libraries were produced by cDNA generation, fragmentation, size selection, PCR amplification, and barcoding.

In order to sequence the libraries, each batch was sent to the Cold Spring Harbor Laboratory sequencing facility. Attachment of library-specific barcodes allowed for pooling of all libraries in the same batch and simultaneously sequencing on an Illumina NextSeq 500 machine to obtain the reads. The FastQC [11] software was used to review read sequencing quality and ensure that low quality sample libraries were excluded in the subsequent processes. A schematic summary of the entire sample-processing pipeline is shown in Fig. 1a. Since most of the acquired samples were of luminal subtype with positive estrogen and/or progesterone receptors status, the analysis exclusively focused on luminal samples for further downstream analysis.

### 2.2 Transcriptional progression of NIS

Inference of gene expression from sequencing data was implemented using a robust pseudoalignment and quantification pipeline, Kallisto [12]. The reads were mapped onto gene isoforms included in the human genome reference annotated by ENSEMBL [13]. This pipeline is supplemented with bootstrap calculations (see Methods) to estimate the uncertainty of each expression value in units of transcript per million (tpm). To ensure accurate estimation and enough gene coverage, we further filtered the samples to those containing at least 10 million mapped reads. From the remaining 28 NIS and 15 epithelial samples, we corrected the normalized gene expression matrix values against prominent batch effects arising from technical experimental artefacts using a linear-based pipeline, ComBat [14, 15].

Given the final expression matrix, we performed principal component analysis (PCA) and projected onto two principal components (PC1 and PC2) with the largest variance as visualized on the 2D plot shown in Fig. 2. Here we label the sample conditions in four different colors. In both NIS (Fig. 2a) and epithelial/cancer (Fig. 2b), prophylactic normal (green) samples appear to be transcriptionally distinct from the tumor positive samples. Intriguingly, ipsi-/contralateral normal (purple) samples (Fig. 2a) are more similar to primary (blue) and metastatic (red) NIS samples, despite looking normal/non-cancerous morphologically. This supports the notion of a cancer field effect, which describes the existence of a zone of influence radiating from cancer cells to surrounding cells through soluble factors such as proinflammatory signals, exosomes, and reactive oxygen species [16, 17].

**Figure 2:**
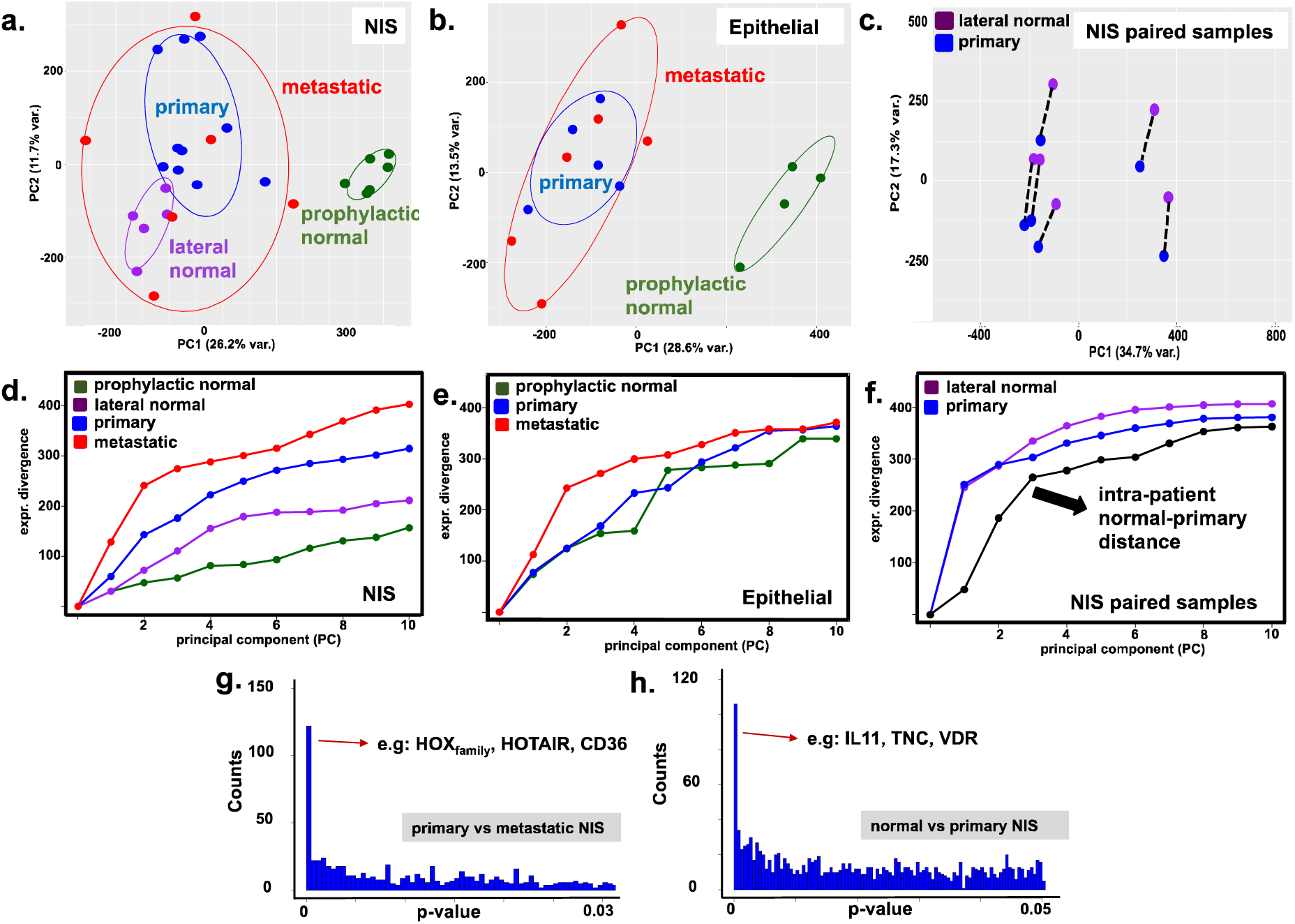
Expression variation and differential analyses. Principal compnent (PC) analyses of genomewide transcription in prophylactic normal, ipsi-/contralateral normal, primary, and metastatic tumor (brain, liver, and bone) of **a)** non-immune stroma (NIS), **b)** epithelial, and **c)** patient-matched (paired lateral normal and primary tumor) NIS samples. Expression divergence (see Methods 4.5) of **d)** NIS, **e)** epithelial, and **f)** paired NIS samples across geographical sites in patients. Distribution of p-values from differential gene expression of NIS estimated for **g)** primary vs metastatic tumor and **h)** prophylactic normal vs primary tumor. For illustration of significant enrichment close to p-value equals zero, the figures are truncated.

In Figure 2a, the primary tumor NIS show significant expression changes compared to ipsi-/contralateral normal tissues. These results support the idea of interactions between NIS cells and epithelial cancer cells in a breast tumor leading to transcriptional modification [18]. Furthermore, the landscape of metastatic samples indicates a wider radial spread across the principal components. Metastatic samples diverge from primary samples nonlinearly due to strong dependence on different metastatic sites. We observed a strong diverging pattern in NIS samples variance (Fig. 2d) across multiple PCs, implying a consistent increase in heterogeneity between individuals from prophylactic normal, to lateral normal, to primary, to metastatic. A similar pattern appears less significantly for epithelial samples as shown in Figure 2e. In conclusion, the transcriptional trajectories of NIS and epithelial samples are not collinear and diverge from normal to primary to metastatic conditions.

Analysis of patient-matched samples was performed to understand how the transcriptional landscape changes from ipsi-/contralateral normal NIS to primary tumor NIS within the same individual. In Figure 2c, we observed that PC2 robustly defines the normal-primary axis of patient-matched NIS samples. Furthermore, we quantified the pair distance from the mean length of dashed lines in Figure 2f. This distance is relatively similar to the standard deviation of samples (see Methods) in each condition, which suggests the degree transcriptional shift from lateral normal NIS to primary tumor NIS is comparable to the heterogeneity of NIS among different individuals.

### 2.3 Differential expressions of exclusive NIS genes

Transcription changes of NIS samples were examined by running a differential analysis on the expression matrix. We applied a conservative approach, Sleuth [19], by taking the gene expressions along with the uncertainties for linear modeling and statistical testing. During the analysis, we incorporated patient, condition, and batch information as variable inputs with the corresponding statistical noise. Differential genes were determined from Wald test results with q-value less than 0.05.

Significant differences between prophylactic normal and primary NIS samples are shown by p-value distribution in Fig. 2h, which depicts some significant genes with p-value less than 0.05. There are approximately 80 differential genes (q-value<0.05) that signify the transcriptional change between the two conditions. Among others, extracellular matrix genes such as *TNC* [20] appear to be significantly downregulated. We also found immune related genes that are involved in inflammatory responses such as *IL11*, *RN7SL1*, *SERPINB9*, and *ALOX5AP* [21–24]. Long noncoding and histone genes were also significantly modified from prophylactic normal to primary conditions. These results indicate signaling interactions of NIS cells with cancer and immune cells in the tumor microenvironment. Furthermore, the significant genes collectively suggest their involvements in pattern formation, tissue development, and embryogenesis.

Comparisons between primary and metastatic tumor NIS were similarly described by significant p-value peak shown in Fig. 2g. We identified 108 differentially expressed genes with q-values less than 0.05, which have no overlap with the differential genes from normal versus primary. Among them, we can find multiple *HOX* genes and *STRIP2* that are important in developmental and morphogenesis [25, 26]. Many growth factors and extracellular genes revealed in the list such as *PGF*, *FGFR2* and *NOG* are significant in metastatic processes [27-29]. We also found *CD36* and *CD14,* which are genes relevant to certain CAF functions such as cell adhesion, collagen binding, and immune response [30, 31]. Some noncoding genes that are well-known cancer markers such as *HOTAIR* and *HAGLR* [32, 33] were downregulated in breast NIS progression. On the other hand, *WNT5B* may be related directly to oncogenesis through the *WNT* signaling pathway [34]. To summarize, we provide the complete list of differential genes in both comparisons and paired analysis in the supplementary material (Table 1, Table 2, and Table 3).

### 2.4. Putative interactions between epithelial/cancer cells and NIS cells

To infer putative crosstalk between cancer cells and NIS cells, the gene expression dataset was cross-referenced to known ligand-receptor pairs. Only genes that were previously identified as significantly differentially regulated between normal versus primary tumor and primary tumor versus metastases were considered. A comparison of gene expression from normal to primary tumor showed the upregulation of the ligand-receptor pairs *GDF5* and activin A receptor (*ACVR1*, *ACVR2A*) in both NIS cells and cancer cells. This pair belongs to the *TGF*-beta super family and is involved in epithelial-mesenchymal transition (EMT) [35, 36].

Further progression from primary tumor to metastatic tumor resulted in increased expression of several more ligand-receptor pairs, including the interaction between cell surface integrin (*ITGA2*) and collagen (*COL7A1*) as well as growth factors (*PGF*, *FGF*) and their respective receptors. Further, direct cell-cell interactions involved in tumorigenesis, angiogenesis, and cancer stem cell generation were also identified (*EPHA* and *EPHB* with *EFNB1*) [37, 38]. Upregulation of these pairs points towards a putative interaction and necessary role between the NIS cells and cancer cells. The list of ligand-receptor pairs with significant NIS q-value less than 0.1 is provided in supplementary material (Table 4 and Table 5).

The role of *p53* in tumor progression has been widely studied and accepted. Mutation and dysregulation of the corresponding *TP53* as a tumor suppressor gene can alter several biological functions that are important in tumor growth such as cell cycle arrest, DNA repair, apoptosis, angiogenesis, and metastasis [39]. Among the genes involved in *p53*-induced metastasis and angiogenesis [40], *Thrombospondin-1* (*THBS1*/*TSP1*) is downregulated in metastatic epithelial/cancer and NIS samples as depicted in Fig. 3a. To understand the crosstalk between cancer and NIS, we estimated the expression change of certain receptors and their ligands that emerged from *p53* downstream pathways. An antiangiogenic receptor *CD36* [41] is also found to be downregulated in metastatic samples, which suggests a possible signal from the epithelial/cancer ligand *THBS1* towards the NIS receptor *CD36* promoting metastasis. Furthermore, opposite interactions are also indicated by the co-regulation between NIS ligand *THBS1* and epithelial/cancer receptors *ITGB3* and *LRP1*. The loss of cell adhesion caused by low expression of *ITGB3* may drive cancer cells to detach [42] and metastasize from the primary site. Together, these results signify the importance of cancer-NIS crosstalk in the progression of breast tumor.

Another way to approximate the interaction between NIS and epithelial/cancer cells is to quantify the mutual information between ligand and receptor pairs, which were listed in a previous study [43]. Mutual information, representing the strength of association for ligand/receptor pairs, was calculated from the t-test significant (p-value<0.1) expression changes between primary and metastatic conditions (see Methods). The advantage of using mutual information rather than any correlation coefficients is its ability to measure association in nonlinear and non-monotonic relationships [44]. We calculated the joint probability by counting the number of ligand-receptor pairs where the primary-metastatic expression changes increased or decreased. In Fig. 3b, we show the mutual information values in bits (corrected for finite number of samples) for random pairs of genes, NIS ligands/epithelial receptors, epithelial ligands/NIS receptors, and overall ligands/receptors. We found that there is a significant increase in the epithelial ligands/NIS receptors mutual information at about 0.021 bits. This is relatively high compared to random and NIS ligands/epithelial receptors which were close to zero. This discrepancy indicates a tendency towards uni-directional communication from epithelial/cancer ligands towards NIS receptors as the tumor progresses from primary to metastatic. Consequently, this supports the interaction between cancer cells and NIS as a part of EMT mechanism and the concept of cancer field effect. The list of ligand-receptor pairs with significant p-value less than 0.1 from primary to metastatic is provided in supplementary material (Table 6 and Table 7).

**Figure 3.**
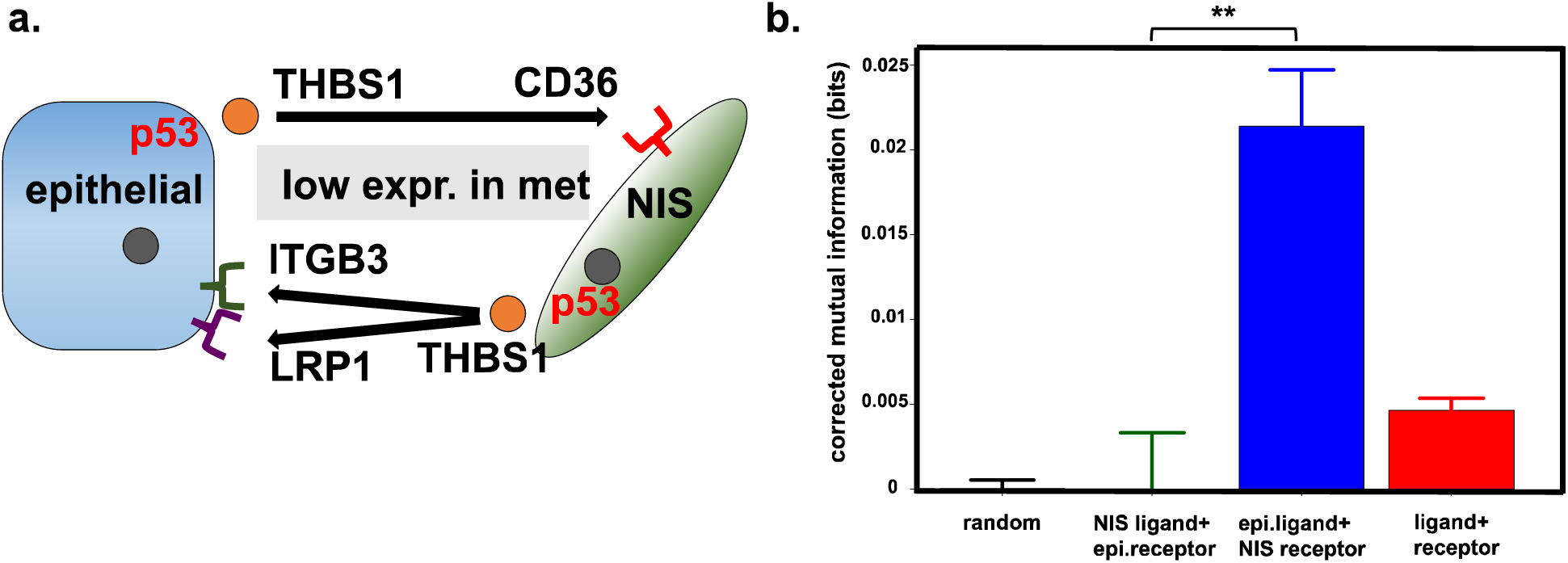
Inference of cancer and non-immune stroma (NIS) crosstalk from RNA-seq data. **a).** Crosstalk between epithelial/cancer and NIS cells through *p53* downstream ligand *THBS1* and its receptors (*CD36*, *ITGB3*, and *LRP1*), which are downregulated in metastatic samples. **b)** Mutual information of expression changes with standard error from primary to metastatic between (left to right): random pairs of genes, NIS ligands/epithelial receptors, epithelial ligands/NIS receptors, and ligands/receptors (**p-value < 2×10^−16^).

### 2.5 C2orf88 expression predicts patient survival

To analyze specific NIS progression in a tumor, we filtered the differential gene expressions with a conditional threshold. The estimation of transcriptional noise becomes very relevant when distinguishing expression that is genuinely caused by active transcriptions. Based on the work by Wagner et al [45], we implemented a threshold value of 2 tpm, which represents the baseline for background/noise transcription. Among the differential genes, we prioritized exclusive NIS genes with an average NIS expression above 2 tpm in at least one tissue condition, but lower than 2 tpm for epithelial samples in every tissue condition.

We investigated the functional role of a novel candidate NIS gene in our list of significant genes, namely *C2orf88*. The function of this protein-coding gene is largely unknown in cancer and only reported to be involved in protein kinase A binding and cell-cell interactions [46]. A previous study identified low expression of *C2orf88* in tumor tissue compared to normal breast tissue using a cDNA microarray assay [47]. This study focused exclusively on the bulk tumor, aggregating the epithelial and stromal tissue, thereby losing information about the cellular source of the gene expression. In our differential analysis, *C2orf88* had significant change in expressions from primary NIS to metastatic NIS with (q-value = 0.019). The expression value was decreased by about 6 fold from 4.5 tpm to 0.7 tpm, as shown in Fig. 4a. Despite being expressed moderately in NIS, the expression values stayed under 2 tpm in epithelial, suggesting that the low residual expression might come from a non-active gene transcription. To investigate the clinical impact of NIS gene expression, we analyzed survival data of luminal breast cancer patients. We analyzed the TCGA expression database with 827 breast cancer patients and grouped them into: ‘high’ expression with values above the third quartile, or ‘low’ expression with values below the first quartile. By implementing Kaplan-Meier [48] analysis (see Methods), as shown in Fig. 4b, we found that the high expressor patients provide better prognosis and survival compared to low expressors with a significantly low (p-value = 3.5×10^−3^). The relative level of C2orf88 expression is phenotypically reflected clinical outcome.

**Figure 4:**
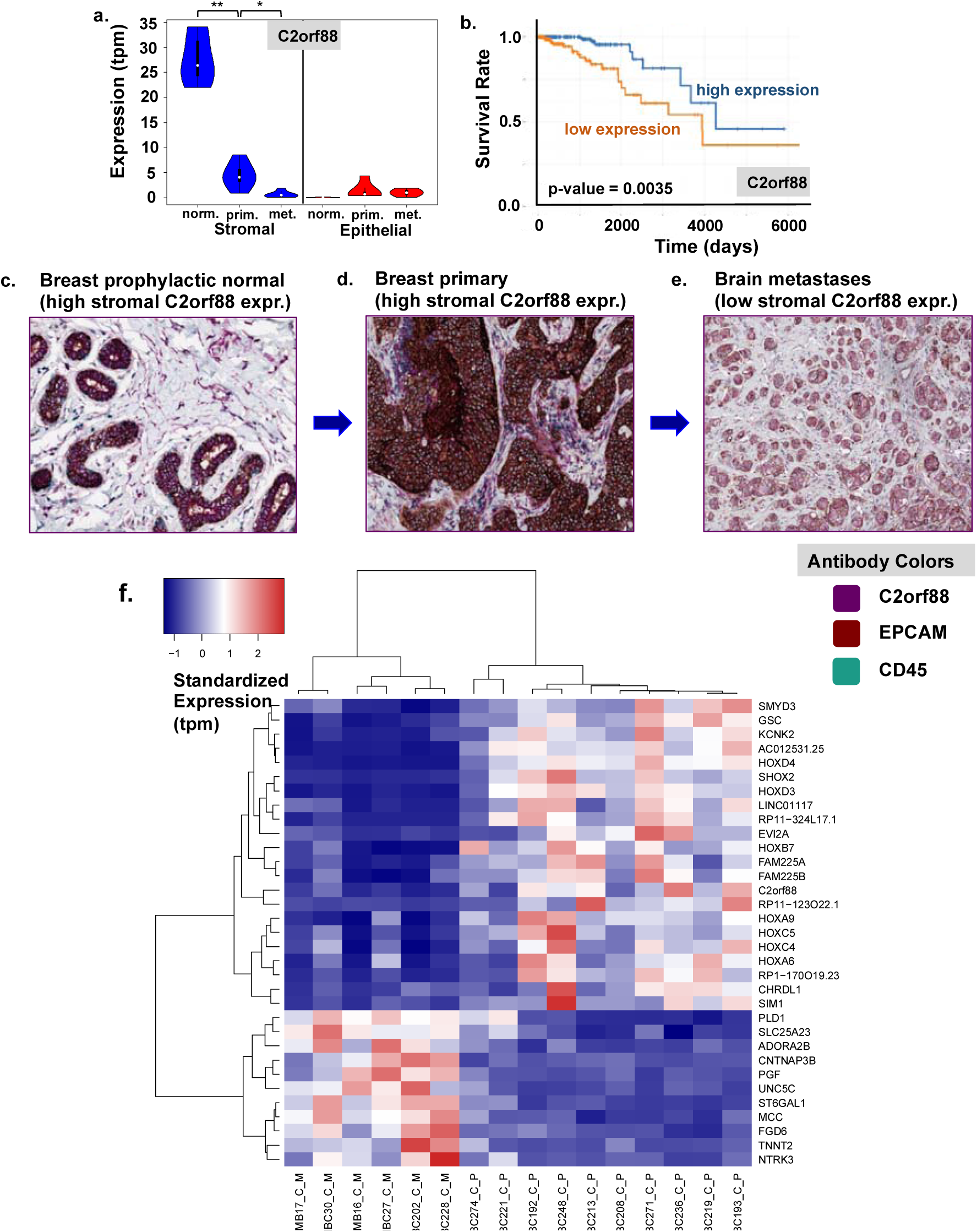
Expression and survival profile of *C2orf88* in non-immune stroma (NIS) and epithelial. **a)** Expression values (tpm) in prophylactic normal, primary, and metastatic tumor samples (**p < 2.13×10^−5^; *p < 6.69×10^−4^). **b)** Survival analysis of luminal breast cancer from TCGA database with Kaplan-Meier plot featuring log-rank p-value between high (>3^rd^ quartile) and low (<1^st^ quartile) expressions. IHC validation (purple: *C2orf88*, brown: *EPCAM*, teal: *CD45*) on **c)** breast prophylactic normal, d) breast primary, and **e)** brain metastases. **f)** Heatmap plot with hierarchical clustering showing significant gene expressions in primary (left group) and metastatic (right group) samples which correlate strongly with *C2orf88* (|ρ|_pearson_ < 0.6)

Representative gene expressions, which correlate with *C2orf88* were clustered and mapped in a heatmap plot (see Methods) of Fig. 4f. Collective genes enrichment estimated the significant role of these differential genes in epithelial-mesenchymal transition (EMT) in primary and metastatic comparisons. These correlative gene expression signatures suggest that NIS cells from primary tumor are expressing EMT-related genes. The EMT genes are further downregulated as the tumor metastasizes and essentially undergoes mesenchymal-epithelial transition (MET) in order to colonize the metastatic site [49].

### 2.6 IHC validates C2orf88 expression levels and NIS heterogeneity

Based on the differential and survival analysis results, we further validated the expression of *C2orf88*, which has shown to be a good candidate for identifying NIS progression in tumors. To investigate *C2orf88* expression change in tumors, we examined the intensity and population of cells that were stained with the antibody in luminal breast tumor tissues. In Fig. 4c, we see distinct *C2orf88* protein expression (purple) in prophylactic normal breast tissue with relatively high intensity in NIS compartments compared to the background counterstaining in blue. We also differentiate between epithelial/cancer and immune cells by staining *EPCAM* (brown) and *CD45* (teal), respectively. A strong *C2orf88* intensity was observed in the NIS of a breast tumor (Fig. 4d). However, this intensity varied significantly among NIS cells, which indicates the existence of heterogeneity within the tumor NIS, which has been a major challenge for many cancer therapies. Furthermore, a consistent expression change was successfully shown by similar results on brain and liver metastatic tissues. Low *C2orf88* expression level in Fig. 4e supports the differential analysis of this gene, which suggests significant down-regulation from primary to metastatic tumor NIS. The expression change in epithelial also indicates EMT as shown in supplementary material (Figure S1). Mesenchymal cells that emerge from dense cancer islands and have lost their epithelial marker (*EPCAM*) have increased *C2orf88* expression. Together, these results reveal a novel gene *C2orf88* expressed in NIS cells that may regulate breast tumor development and progression.

## 3 Discussion

The role of stroma in the progression of breast cancer remains a subject of ongoing research. From expression profiles of NIS cells, previous studies have implied that several genes are involved in promoting tumor growth, invasion, and metastasis. Deep-RNAseq and subsequent analysis on short-term cultured patient tissues allowed us to obtain transcriptional landscapes of NIS from primary and metastatic tumors as well as non-cancerous breast tissues. Accurate computational methods for expression quantification and differential analysis were implemented to capture the expression changes between tumor conditions and then correlate it with patient survival.

Here we show that NIS cells in macroscopically normal breast tissues, adjacent to the tumor, are transcriptionally different from non-cancerous breast tissues obtained from prophylactic mastectomy patients with *BRCA* mutation. These findings support the concept of a cancer field effect [16]. NIS from primary and metastatic tumors undergo further significant expression changes from adjacent normal tissues, involving different sets of genes. As breast cancer progresses into metastatic sites, the transcriptional landscapes of both cancer and NIS cells diverge due to the distinct environment within the different tissue sites. As shown in Fig. 3b, our novel crosstalk analysis suggests that metastatic cancer cells actively modulate NIS cells to condition its microenvironment for survival. These results lend further support to the idea that breast cancers dynamically interact with their microenvironment.

In summary, this work unravels significant transcriptional changes of NIS between different tumor conditions in luminal breast cancer. Several coding and long noncoding genes that are known to be involved in embryonic morphogenesis, immune responses, and epithelial-mesenchymal transition, such as *HOX* family, Histone family, *C2orf88*, *ALOX5AP*, *IL11*, *CD36*, *TNC*, *VDR*, and *HOTAIR* were found to be differentially expressed. Supplemented by TCGA survival analysis, expression of the downregulated *C2orf88* gene was validated with IHC staining. These results provide a new understanding of NIS transcriptional progression and its corresponding genes involved in the evolution of breast cancer, suggesting potential therapies to directly target the tumor NIS.

## 4 Materials and Methods

### 4.1 Tissue and culturing

Normal breast tissue from *BRCA*+ patients undergoing a prophylactic mastectomy, primary breast tumor, and tumors from metastatic sites were obtained from patients treated at City of Hope according to guidelines approved by City of Hope Institutional Review Board. Tissues were dissected into small pieces (1-2 mm diameter), and 5-6 pieces per well were cultured in a 6-well culture plate (Fisher Scientific). For up to two weeks tissue pieces were cultured in Advanced Dulbecco’s modified Eagle’s medium (DMEM) supplemented with 6.5% FBS, 0.5% MEGS and 1% penicillin streptomycin. If longer culturing was required to grow sufficient cells, media was switched to Advanced DMEM with 10% FBS and 1% penicillin streptomycin for an additional week.

### 4.2 Cell sorting and RNA extraction

After the culture well reached confluency, cells were trypsinzed (0.25%) and stained for fluorescent cells sorting using BD FACSARIA III. The cells were stained labeled with *CD45* PE (Leukocytes) to minimize immune contamination (Biolegend #BL304008, 1:200), *EPCAM*/*CD326* APC (Biolegend #324208, 1:100) *CD44* FITC (Biolegend #BL338804, 1:200) for 15 min at room temperature. All cells were gated for *CD45*-expression and then further gated for *CD44*+ (fibroblasts) and *EPCAM*+ (cancer cell) expression. Cells were centrifuged at 300g for 5 min and RNA extracted using Qiagen RNeasy Plus Micro kit (#74034) according to manufacturer’s protocol. Illumina sequencer compatible library was generated with Kapa mRNA Hyper kit (Roche, Basel Switzerland) for polyA library preparation according to manufacturer’s protocol.

### 4.3 Deep RNA sequencing

Libraries were pooled with equal moles based on qubit mass measurements and expected fragment length. The libraries were sequenced on NextSeq 500 using a 75-cycle v2 SBS kit. There was a 5% PhiX used as a control sample and the sequencing was done as a single-end 76 length with an index read. Reads were demultiplexed by barcode via the bcl2fastq2 tool.

### 4.4 Data and source code availablity

The raw sequence fastq data have been deposited in Sequence Read Archive (SRA) repository with accession code: SRP157590. The source code written in R for the bulk RNA-Seq analyses are available on https://github.com/AtwalLab/BulkRNAseq

### 4.5 Pseudoalignment and quantification

The RNA-seq sequence reads were pseudoaligned to coding and long noncoding regions in ENSEMBL Human genome (GRCh38 rel85) using an open-source package, Kallisto. The gene expression was quantified by “kallisto quant” with option attachments as follows: single end reads, first read reverse, bias correction, seed 42, fragment length 200 ± 20 bp, and bootstraps 100, and number of threads 4.

Principal component analysis and plotting was performed with the R function, “prcomp”, and package, ggbiplot. Expression divergences were calculated by the cumulative sum of the *i*-th PC eigenvalues (***e_i_**)* that correspond to the variance, as follows:

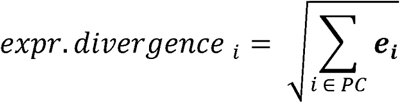

The eigenvalues correspond to the variance of the samples in principal components space.

### 4.6 Differential analysis with batch correction and heatmap clustering

Differentially expressed genes were estimated using an open-source package Sleuth. We fit the tissue condition, patient ID, and batch as covariates in the full model and only set the batch in the reduced model. Aggregation was made on ENSEMBL gene ID and we implemented Wald test on tissue conditions for the full model. Similar settings were applied for batch correction pipeline using an open-source package, ComBat on log2 normalized expression matrix. Heatmap visualization of the expressions was generated using heatmap3 package with default setting, as follows: standardized expression (tpm) across samples, hierarchical clustering, complete linkage, and distance (1 - pearson correlation).

### 4.7 Mutual information and survival analysis

The mutual information (I), in bits, between ligands expression change, L, and their associated receptors R expression change from primary to metastatic was calculated as:

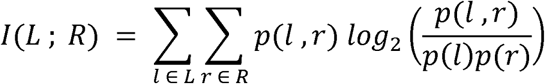

The joint probability was calculated by counting the number of ligand-receptor pairs that were transcriptionally upregulated or downregulated. In order to correct for biases in our MI estimate arising from our limited sample sizes, we then applied a bootstrapping based finite-sampling correction previously described [44, 50]. The survival analysis was based on Kaplan-Meier plots from an open-source package, RTCGA. The TCGA expression database was filtered for luminal breast cancer patients and grouped into patients with expressions above 3^rd^ quartile and below 1^st^ quartile. The p-value was estimated by log-rank test between the two groups.

### 4.8 Immunohistochemistry staining

Triple chromogenic IHC was performed on Ventana Discovery Ultra platform at CSHL Histology Core, with OmniMap HRP, ChromoMap DAB, Discovery Purple, and Discovery Teal detection systems according to manufacturer’s protocols (Roche). Antibodies: *C2orf88* (Thermofisher, dilution 1:50), *CD45* (Abcam, 1:500), and *EPCAM* (Ptglab, 1:200)

## Acknowledgements

We would like to thank the sample acquisition team at City of Hope and Histology Core at Cold Spring Harbor Laboratory with NIH support grant 5P30CA045508. This work was supported by aConvergence Award from Stand Up To Cancer and Breast Cancer Research Foundation (BCRF). RU and AB were funded by postdoctoral fellowships from The V Foundation.

## Author Contributions

R.U.: computational analysis, interpretation, IHC design, and manuscript writing; A.B.: data collection, library preparation, and interpretation; N.S.: computational analysis; Y.J.: computational analysis; E.A.: sequencing; Q.G.: IHC staining; Y.H.: data collection and library preparation; S.G.: data collection and library preparation; P.L.: conception, interpretation, and manuscript writing; G.A.: conception, interpretation, and manuscript writing

